# Dopaminergic innervation at the central nucleus of the amygdala reveals distinct topographically and functionally segregated regions

**DOI:** 10.1101/2022.09.22.508929

**Authors:** Eric Casey, María Elena Avale, Alexxai Kravitz, Marcelo Rubinstein

## Abstract

The central nucleus of the amygdala (CeA) is involved in the expression of fear and has been implicated in several anxiety disorders. Anatomically, it is divided in a medial (CeM), a lateral (CeL) and a capsular division (CeC). The CeA is densely innervated by dopaminergic projections that originate in the ventral periaqueductal gray/dorsal raphe (vPAG/DR) and the ventral tegmental area/substantia nigra compacta (VTA/SNc). However, it is unknown if DA exerts a homogenous control over the CeA or, conversely, if different parts of the CeA are regulated in a distinct manner. Here, we performed a neuroanatomical and functional analysis of the mouse CeA and revealed that dopaminergic innervations arriving from the PAG/DR and the VTA/SNc constitute distinct, non-overlapping pathways that differ in their expression of dopamine transporter. By quantifying the distribution of DAergic fibers and the origin of the innervation, we identified two regions in the CeL: a frontal region innervated by both the VTA/SNc and the vPAG/DR, and a caudal region innervated only by the vPAG/DR; and three regions in the CeC: a fronto-dorsal region innervated only by the VTA/SNc, a fronto-ventral region with sparse DAergic innervation, and a caudal region with low innervation from the vPAG/DR. In addition, we found that each region displays a unique pattern of cFos activation after the administration of cocaine, SKF 38393, quinpirole or haloperidol, demonstrating that the regions identified here are functionally distinct from each other. In summary, this analysis reveals unique properties of the DAergic pathways innervating the CeA, and distinguishes six topographically segregated and functionally distinct regions in the CeA. This unanticipated level of functional heterogeneity calls for more precise anatomical specificity in future functional studies of the CeA.

## Introduction

The central nucleus of the amygdala (CeA) is a neural hub that integrates environmental and internal cues related to fear and anxiety and elicits conditioned behavioral responses. Studies in the CeA have gained considerable interest given its involvement in several pathological conditions such as anxiety disorders and drug addiction [1–4]. The CeA is divided in three anatomical portions defined as medial (CeM), lateral (CeL) and capsular (CeC) [5], each containing distinctive neuronal populations characterized by different molecular markers, connectivity and functions [6,7].

The CeA contains neurons expressing either dopamine D1 (D1R) or D2 receptors (D2R) [7–9] and is densely innervated by dopaminergic fibers [9,10]. Functional studies have implicated dopamine (DA) in the CeA with a variety of behaviors including impulse control [11], fear conditioning [8,9,12–14], fear expression [13,15] and defensive behaviors [16,17]. Despite that, the DAergic circuits that regulate the CeA have been poorly studied in comparison with striatal pathways, and their anatomy and connectivity remain mostly unknown.

DAergic innervation of the CeA originates mainly in neurons located in the ventral periaqueductal gray and dorsal raphe (vPAG/DR), as well as neurons present in the ventral tegmental area and the substantia nigra compacta (VTA /SNc), respectively [18–20]. Although the distance between inputs suggests that vPAG/DR->CeA and VTA/SNc->CeA circuits constitute independent pathways, it is unclear whether innervation arriving from each one converges into the same CeA targets or whether they regulate different parts of the CeA. This constitutes a critical gap because the CeA contains a high diversity of neuronal types with specific functions and topographical locations [21].

The goal of this work was to evaluate if vPAG/DR->CeA and VTA/SNc->CeA DAergic pathways innervate different topographical regions of the CeA, and if those regions are functionally different in terms of DAergic neurotransmission. We performed a detailed neuroanatomical characterization of DAergic inputs at the CeA and revealed that vPAG/DR and VTA/SNc projections to the CeA are mostly non-overlapping pathways expressing different levels of the DA transporter (DAT). A careful analysis of DAergic innervation at the CeA allowed us to differentiate a fronto-dorsal, fronto-ventral and caudal region in the CeC, and a frontal and caudal region in the CeL. Finally, using an *in vivo* pharmacological approach coupled to the detection of *c-Fos* expression in the CeA, we demonstrated that these regions are functionally distinct from each other.

## Results

### DAergic innervations at the CeA

We first analyzed the distribution of DAergic fibers innervating the CeA and determined their relative density across the fronto-caudal extension of each division by tyrosine hydroxylase (TH) immunofluorescence. We found that the most densely innervated area of the CeA is the CeL, while the CeM showed only moderate innervation, and the CeC exhibited an even lower density of TH immunoreactive fibers (**Fig. 1A** and **1B**). While we did not detect a fronto-caudal gradient of DAergic innervation in any division of the CeA (**Fig. 1B**), we found a decreasing dorso-ventral gradient at frontal regions of the CeC (anterior to −1.22mm, **Fig. 1C**). Further analysis of fronto-dorsal and fronto-ventral regions of the CeC evidenced a fronto-caudal gradient only in the CeC (**Supp. Fig. 1**). By dividing the CeA divisions according to the gradients described above, we found significant differences in the density of DAergic innervation at the different regions (**Fig. 1D**). Consistently, we found a similar distribution of fibers expressing the red fluorescent marker *tdTomato* in double transgenic mice carrying a *Dat*^*IRES-Cre*^ *knockin* allele [22] and the Cre-inducible reporter gene Ai14 [23], confirming the DAergic identity of these projections (**Supp. Fig. 2**).

**Figure 1.**
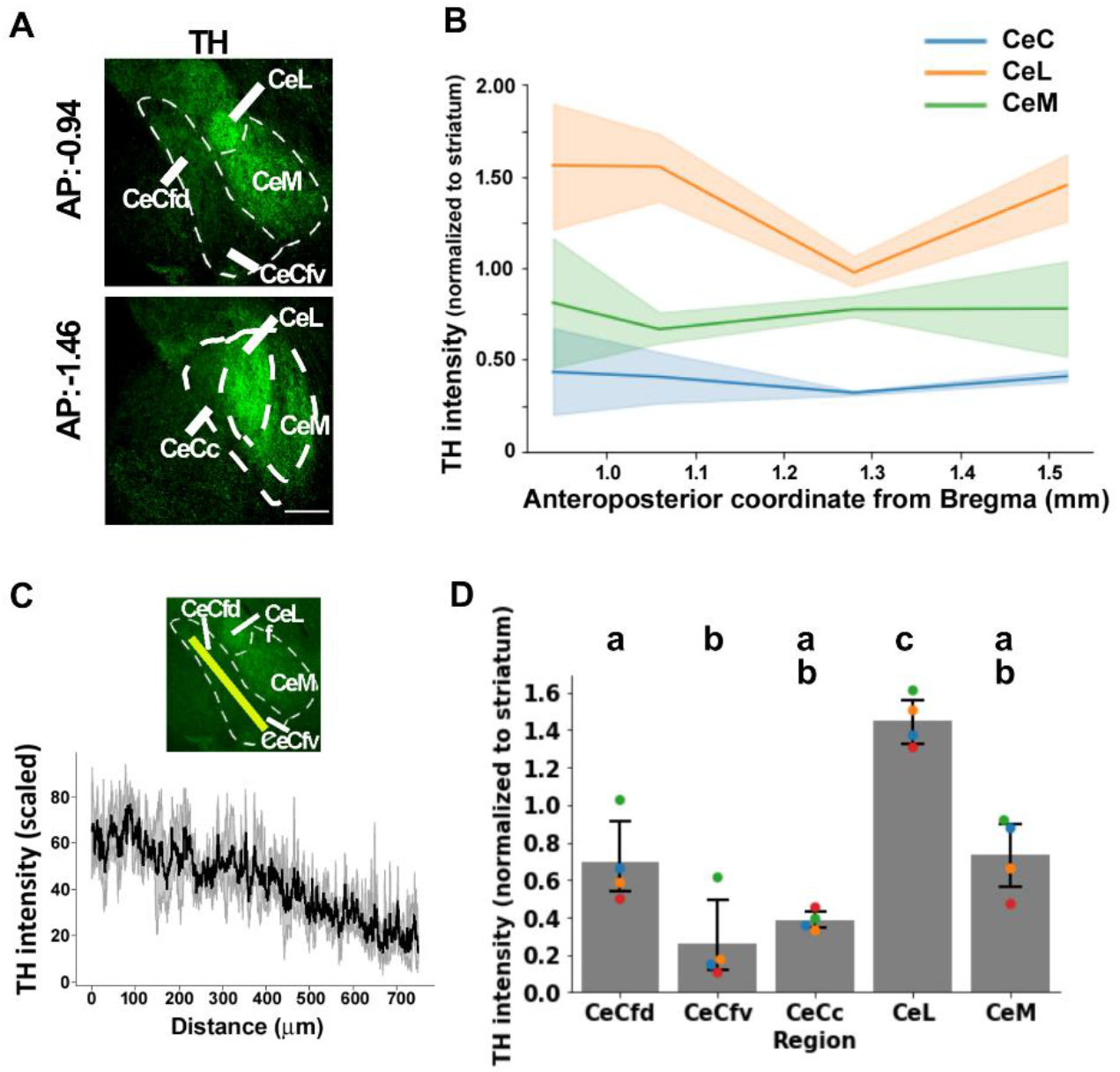
Distribution of DAergic fibers in the CeA. **A)** Representative histology of the CeA of a *wild-type* mouse immunostained against TH. Upper row: frontal part of the CeA, from AP:-0.82 to AP:-1.06 mm (the example corresponds to AP:-0.94 mm); bottom row: caudal part of the CeA, from AP:-1.22 to AP:-1.70 (the example corresponds to AP:-1.46 mm). **B)** Quantification of TH intensity across the fronto-caudal axis, for each division. Values are normalized to intensity in the dorsal striatum of the same coronal section. Significances inferred through Akaike information criterion (AIC) of nested models indicated significant effect of division, but not significant effect of fronto-caudal coordinate nor interaction (“Null model”, AIC= 88.3; “TH ∼ Division” model, AIC= 20.6; “TH ∼ Division + Coordinate” model, AIC= 21.6; “TH ∼ Division x Coordinate” model (with interaction), AIC= 24.4). **C)** Dorso-ventral gradient of DAergic fibers in the frontal CeC. Top: representative histology of the frontal part of the CeA immunostained against TH. Bottom: quantification of fluorescence intensity (in bits) through the transect marked by a yellow line in the top figure, the average values ± standard deviation corresponding to 4 slices from 3 different mice are shown. Linear regression indicated a significant negative slope; linear mixed model, Wald’s test: Intercept, value= 67.1, p<0.001; Slope, value= −0.065, p<0.001. **D)** Quantification of intensity of TH immunofluorescence in different regions of the CeA; each dot indicates individual data, bars and errors indicate the mean ± 95% confidence interval for each region. Repeated measures ANOVA, p<0.0001. Post-hoc Bonferroni test is indicated; same letter indicates not-significant differences (p > 0.05) and different letters indicate significant differences (p < 0.05). CeL, lateral division of the central amygdala; CeCfd, capsular division of the central amygdala, frontodorsal part; CeCfv, capsular division of the central amygdala, frontoventral part; CeCc, capsular division of the central amygdala, caudal part; CeM, medial division of the central amygdala. Scale bars: 200 µm.

**Figure 2.**
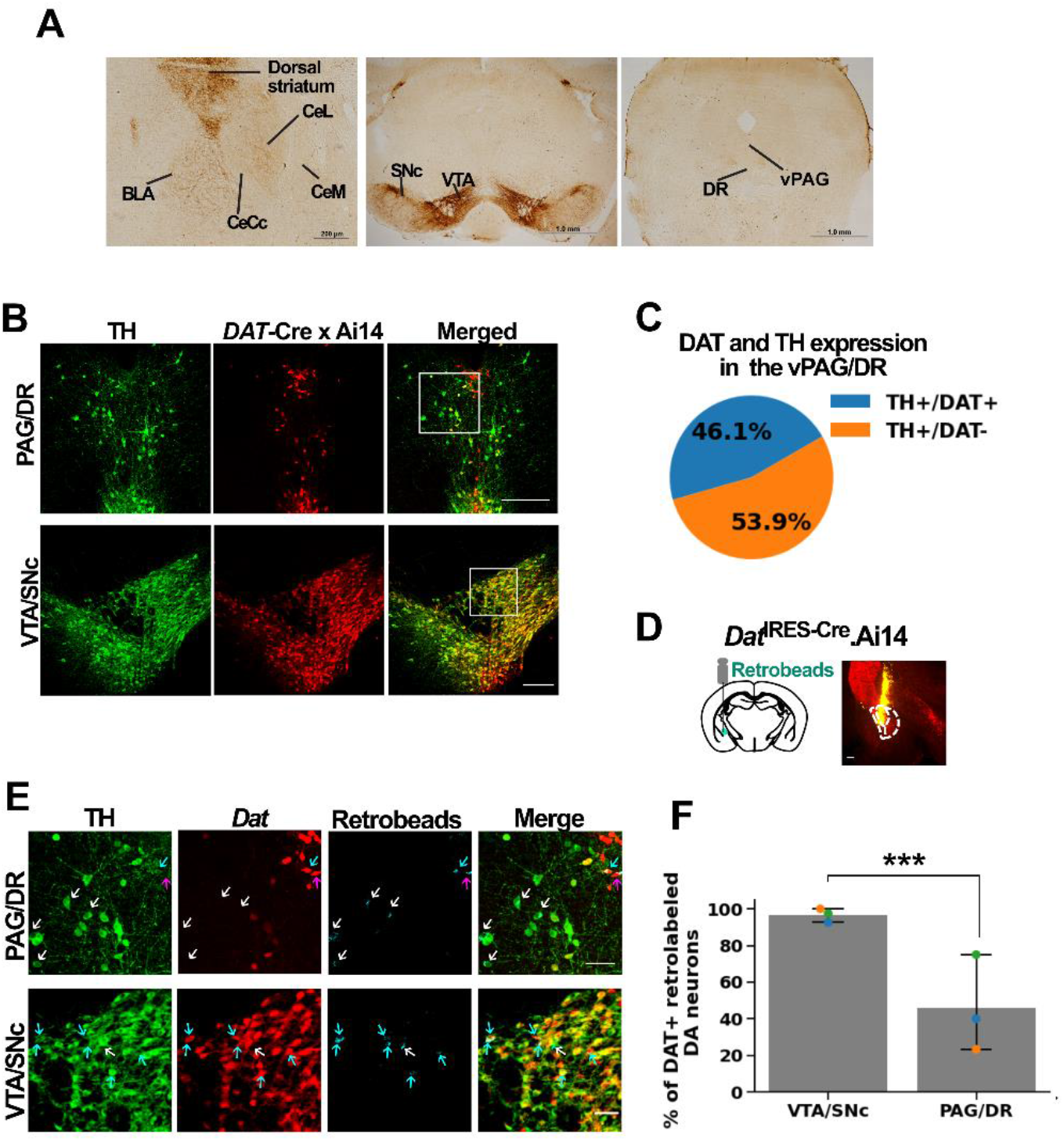
vPAG/DR DAergic neurons display limited expression of DAT. **A)** Immunohistochemistry against DAT on coronal sections containing the CeA (left), the VTA/SNc (center) and the vPAG/DR (right). **B-C)** Colocalization of *Dat* (*tdTomato*) and TH (immunofluorescence) was analyzed in *Dat*^*+/IRES-Cre*^.Ai14 mice. **B)** Representative coronal sections. **C)** Quantification; values represent the average percentage of neurons coexpressing *Dat* and TH or TH alone (n=3 mice). **D-F)** Retrobeads were injected in the CeA of *Dat*^*+/IRES-Cre*^.Ai14 mice and the colocalization of *Dat* (*tdTomato*) and TH (immunofluorescence) was analyzed in the retrolabeled neurons of the vPAG/DR and VTA/SNc. **D)** Scheme of the surgery and representative histology at the injection site. **E)** Magnifications of the squared areas in **B**, showing retro-labeled neurons (cyan). White arrows indicate examples of retrolabeled TH+/*Dat*-neurons; light blue arrows indicate retrolabeled TH+/*Dat*+ neurons; pink arrows indicate retrolabeled TH-/*Dat*+ neurons. **F)** Percentage of *Dat*+ neurons from all retrolabeled and TH+ neurons, in the VTA and in the PAG/DR. Bootstrap with 10000 replicates, p<0.0001, n=3 mice). Scale bars: 200 µm (**A, left** and **B**); 1.0 mm (**A, center** and **right**); 50 µm (**E**).

### vPAG/DR DAergic neurons display limited expression of DAT

In contrast to the intense TH immunolabeling found in the CeA, immunohistochemistry performed using an antibody raised against the DA transporter (DAT) only showed weak signal in the CeA (**Fig. 2A, left**), suggesting that DAT levels in neurons innervating the CeA is relatively low. Then, we evaluated the presence of DAT in the two main DAergic inputs of the CeA and found that while DAT immunohistochemistry strongly labeled neurons from the VTA/SNc (**Fig. 2A middle)**, it was almost undetectable in neurons from the vPAG/DR (**Fig. 2A right**). To further confirm this result with a more sensitive approach, we evaluated the colocalization of DAT, using *tdTomato* as a reporter in *Dat*^*+/IRES-Cre*^.Ai14 mice; and TH, using green immunofluorescence. We found that only half (49.1 ± 4.1 %) of TH immunolabeled neurons in the vPAG/DR also expressed *Dat* (**Fig. 2B-C**). By injecting fluorescent retrobeads (Lumafluor) in the CeA of those mice (**Fig. 2D**), we found that both *Dat*+ and *Dat*-neurons were retrolabeled with retrobeads in the vPAG/DR, demostrating that both types of DAergic neurons project to the CeA (**Fig. 2E**). Furthermore, while almost all neurons projecting from the VTA to the CeA coexpresed *Dat* and *Th*, the amount of retrolabeled TH+ neurons expressing *Dat* in the vPAG/DR was significantly lower (**Fig. 2E-F**), demonstrating that CeA inputs from the VTA and the vPAG/DR differ substantially in their expression of DAT.

### Distinct DAergic innervations from the vPAG/DR and VTA/SNc to the CeA

The CeA receives DAergic inputs arriving from the VTA/SNc and the vPAG/DR [18–20], but it was not clear whether both pathways converge in the same areas or rather assemble different circuits that control specific subdivisions of the CeA. To evaluate these two alternatives, we performed a double immunofluorescence study using antibodies against DAT and TH. We reasoned that, if innervation arriving from the VTA/SNc or the vPAG/DR were topographically segregated, we would detect a heterogeneous immunofluorescence distribution of DAT/TH signal ratios in the different regions of the CeA. Conversely, if both pathways were to converge into the different CeA regions, we would observe a homogeneous distribution of DAT/TH signal at the target sites. In agreement with the results reported above, inputs from the VTA/SNc showed strong labeling for both TH and DAT, while inputs from the vPAG/DR showed undetectable DAT (**Fig. 3A**). In support of the hypothesis of segregated circuits, we found that DAT/TH ratios were highly heterogeneous in the CeA: the frontal CeC has the highest DAT/TH ratio, while the caudal CeC, caudal CeL and CeM have considerably lower DAT/TH ratios. In turn, the frontal CeL showed intermediate DAT/TH ratios (**Fig. 3B** and **Suppl. Table 1**). To further confirm these results, we labeled *Dat*-expressing neurons with a red fluorescent marker by stereotaxic injections of adeno-associated viral (AAV) particles expressing a Cre-inducible *mCherry* reporter into the vPAG/DR or VTA/SNc of *Dat*^*+/IRES-Cre*^ mice (**Fig. 3C**). Analysis of *mCherry* fluorescent fibers arriving at the CeA from the vPAG/DR or the VTA/SNc revealed two different innervation patterns. The Cre-inducible *mCherry* AAV particles injected into the VTA/SNc of *Dat*^*+/IRES-Cre*^ mice labeled fibers in the frontal part of the CeC (AP:-0.82 mm to AP:-1.06 mm) (**Fig. 3D, top left**), and only a sparse array of fibers in the caudal part of the CeL/CeC (AP:-1.22 mm to AP:-1.70 mm) and the CeM (**Fig. 3D, top right**). In contrast, injections of the same AAV into the vPAG/DR of *Dat*^*+/IRES-Cre*^ mice labeled a dense array of fibers innervating the caudal portion of the CeL (**Fig. 3D, bottom right**) and, to a lesser extent the CeM, but not the frontal CeC (**Fig. 3D, bottom left**). In turn, the frontal part of the CeL (AP:-0.82 mm to AP:-1.06 mm) was strongly labeled with fibers projecting from both the VTA/SNc and vPAG/DR (**Fig. 3D, top and bottom left**). These results are also consistent with data from the Allen Mouse Brain Connectivity Atlas [24] showing *Dat* and *Th* expressing projections from the VTA/SNc and vPAG/DR to the CeA (**Suppl. Fig. 1**). Together, these results reveal the existence of two distinct DAergic pathways innervating the CeA: 1) PAG/DR neurons projecting mainly to the caudal CeL and CeM that display low DAT levels (**Fig. 3E**) and, 2) VTA/SNc neurons projecting mainly to the frontodorsal CeC expressing high DAT levels (**Fig. 3E**). Interestingly, the unique region of the CeA in which both pathways seem to overlap is the frontal part of the CeL (**Fig. 3E**).

**Figure 3.**
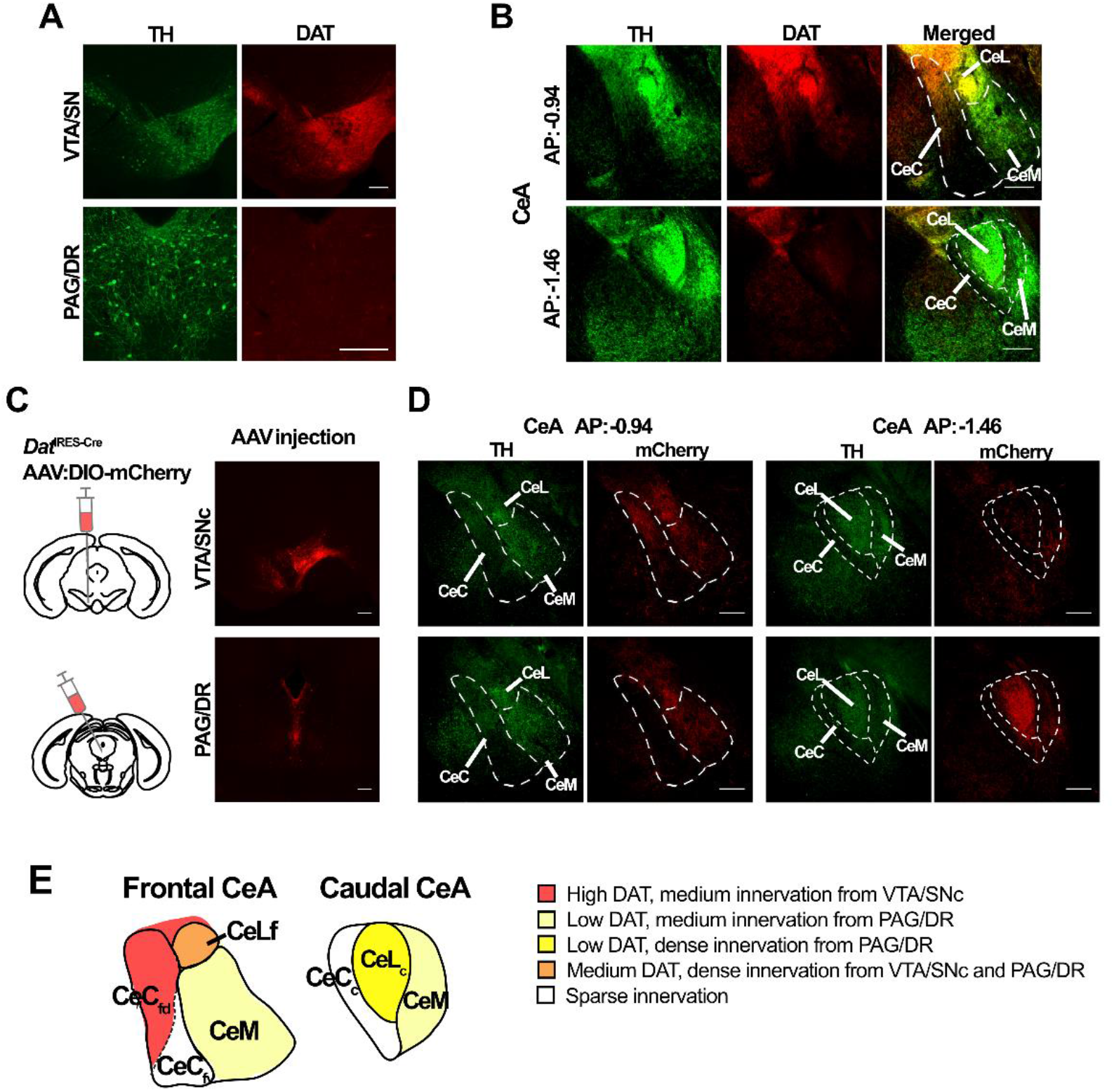
DAergic inputs from the PAG/DR and VTA/SNc display distinct distributions in the CeA. **A-B)** Immunofluorescence against TH (green) and DAT (red) in the VTA/SNc (**A, top**), the PAG/DR (**A, bottom**) and the CeA **(B). C)** Representative coronal sections of a *Dat*^+/IRES-Cre^ mouse brain injected with Cre-inducible ChR2-mCherry in the VTA/SNc (top) or the vPAG/DR (bottom), and **D)** resulting labeling of fibers (red) in frontal (left) and caudal (right) CeA and immunofluorescence against TH (green). **E**) Schematic depicting the distribution of DAergic fibers of each pathway in the CeA. The frontodorsal (fd) and frontoventral (fv) parts of the CeC are separated by dashed lines to highlight their different DAergic innervation content. CeLf, lateral division of the CeA, frontal part; CeLc, lateral division of the CeA, caudal part; CeCfd, capsular division of the CeA, frontodorsal part; CeCfv, capsular division of the CeA, frontoventral part; CeCc, capsular division of the CeA, caudal part; CeM, medial division of the CeA. Scale bars: 200 µm.

### CeA regions defined by their type of innervation are functionally different

Our analysis of the DAergic innervation allowed us to differentiate between a fronto-dorsal, a fronto-ventral and a caudal region in the CeC (CeCdf, CeCfv and CeCc, respectively) and a frontal and a caudal region in the CeL (CeLf and CeLc, respectively) (**Fig. 3E**). Then, we decided to evaluate whether those topographical variations in the density of DAergic inputs involve differential processing of signals in the CeA. We reasoned that if those regions were functionally different, they would show distinct patterns of activity after specific pharmacological manipulations. To this end, we assessed the effects of DAergic compounds injected to live mice in the activation of CeA neurons, as determined by the expression of the immediate early gene *c-Fos*. The indirect DA agonist cocaine (20 mg/kg, i.p.) increased the number of c-FOS+ nuclei only in the caudal part of the CeL and the caudal and frontoventral subdivisions of the CeC **(Fig. 4A-B**). Similarly, the D2R-like agonist quinpirole (1 mg/kg, i.p.) increased the amount of c-FOS+ nuclei exclusively in the caudal part of the CeL and the CeC, but not in the CeM or any other frontal region of the CeA (**Fig. 4C-D**). In contrast, the D2R-like antagonist haloperidol (0.3 mg/kg, i.p.) increased c-FOS immunoreactivity only in frontal CeL neurons (**Fig. 4C-D**). Finally, the D1R agonist SKF 38393 (4 mg/kg, i.p.) strongly increased the number of c-FOS+ nuclei in the fronto-ventral part of the CeC and, to a lesser extent, in the fronto-dorsal CeC, caudal CeC and caudal CeL (**Fig. 4C-D**). Analysis of c-Fos expression across the antero-posterior axis of the CeA further highlighted the existing differences in regions of the same division (**Fig. 4E**).

**Figure 4.**
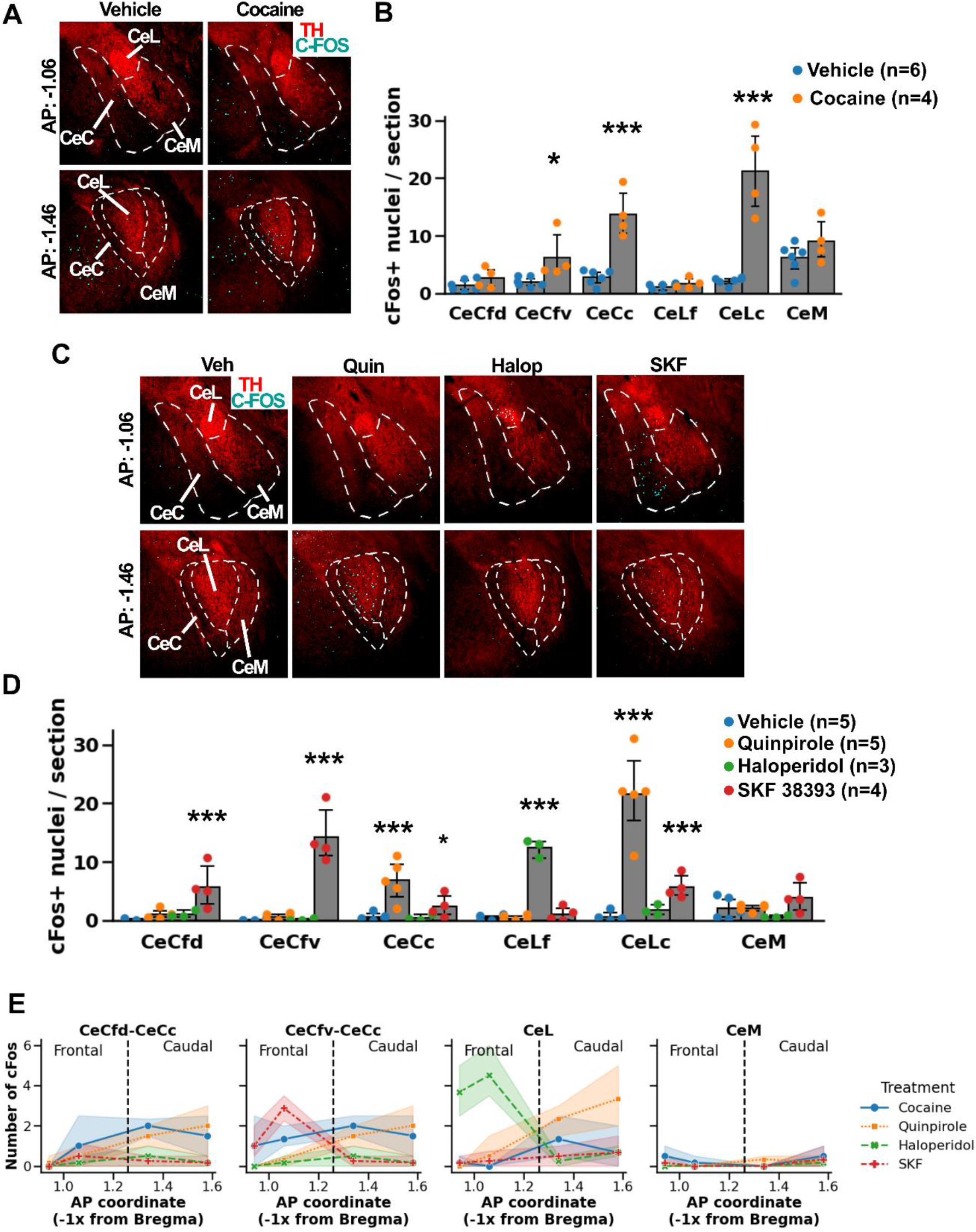
Effect of DAergic drugs on neuronal activity of the CeA. **A-B)** c-*Fos* activation in the CeA induced by saline (vehicle) or cocaine. **A)** Representative frontal (top) and caudal (bottom) CeA sections showing immunofluorescence for TH (red) and c-FOS (cyan). **B)** Quantification of c-FOS positive nuclei in the various CeA regions in mice receiving saline or cocaine. Generalized linear mixed model with negative Binomial distribution (link: log); likelihood-ratio test, Region*Drug: x^2^(5) = 32.1, p<0.0001; post hoc Scheffé test, statistical differences between drugs for each division are shown. **C-D)** c-FOS expression in the CeA induced by saline (Veh), quinpirole (Quin), haloperidol (Halop) or SKF 38393 i.p. injections. **C)** Representative frontal (top) and caudal (bottom) CeA sections showing immunofluorescence for TH (red) and c-FOS (cyan). **D)** Quantification of c-FOS positive nuclei. Two-way generalized linear mixed model with Poisson distribution (link: log); likelihood-ratio test, Region*Drug p < 0.0001; post hoc Scheffé test, statistical differences between drugs for each division are shown. **E)** Number of c-Fos expressing cells per sampling area (shown in **Fig. 5A**) across the fronto-caudal axis after injections of each drug, for each division (2 plots for the CeC highlight differences between Cefd and CeCfv). Each value indicates the mean ± 95% confidence interval across samples and mice. *p<0.05, **p<0.01, ***p<0.001.

While these results imply that distinct CeA regions are differentially activated by DAergic compounds, the identification of functional distinct areas using a pre-conceived anatomical compartmentalization of the CeA could be somewhat biased. For example, if there were different patterns of activation in the CeM across its fronto-caudal axis, this method would not detect them simply because we did not delimit different regions in the CeM (**Fig. 3E**). To overcome this limitation, we sought to further analyze these responses using an unsupervised leaning method that avoids pre-conceived region delimitations bias. To this end, we randomly selected two areas per region in each of the four coronal sections collected in equivalent antero-posterior coordinates for each tested mouse. This procedure allowed us to analyze results in four samples per region except the CeM, which had eight samples (**Fig. 5A**). Then, we calculated in every sample the level of c-Fos expression induced by each drug, and multivariate responses (c-Fos activity for every drug) corresponding to each sample were analyzed following a principal component analysis (PCA). We found that the three first principal components (PC1, PC2 and PC3) explained over 90 % of the variance of the data, mostly influenced by haloperidol, quinpirole and SKF 38393, respectively (**Suppl. Table 2**). Scatterplots of principal component values revealed that samples corresponding to regions identified by their DAergic innervation tended to group together: CeLf samples grouped in a cluster with the highest PC1 values, CeCc and CeLc samples clustered with the highest PC2 values, and CeCfv samples clustered with the highest PC3 values. Finally, CeM and CeCfd samples mostly clustered together at the lowest values of each component (**Fig. 5B-C**). Therefore, the use of an unsupervised learning algorithm to analyze drug-induced c-Fos activation along the various CeA regions is coincidental with those identified based on the distinct patterns of DAergic innervation, supporting the functional identity of the regions delimited based on their innervation. Altogether, these results reveal that each CeA region displays a unique pattern of activation when treated with pharmacological compounds that affect DAergic neurotransmission: the frontal part of the CeL was activated exclusively by D2R-blockade, the fronto-dorsal part of the CeC was activated exclusively by D1R stimulation (although this effect was subtle and for this reason it grouped with the CeM in the PCA), the fronto-ventral CeC was activated by D1R stimulation and DAT blockade, and the caudal CeC and caudal CeL were activated by D1R stimulation, DAT blockade and D2R stimulation. Differently, the CeM did not show any increase in c-FOS following any of the pharmacological manipulations used in this study, compared with saline injections (**Fig. 5D**). In summary, these results demonstrate that the topographically segregated pattern of DAergic innervation of the CeA delimits sub-regions with distinct functional properties.

**Figure 5.**
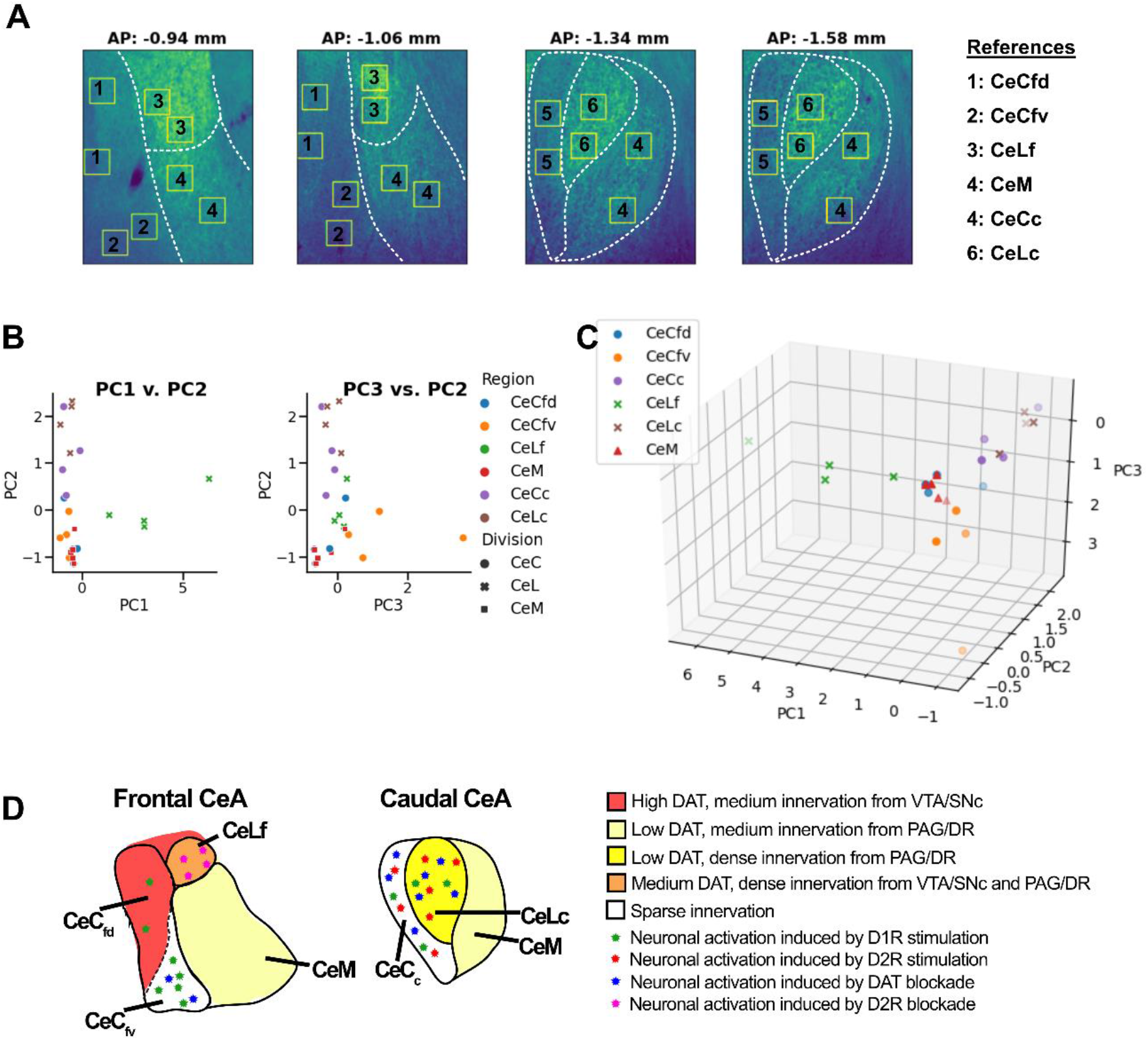
Functional neuroanatomy of DAergic neurotransmission in the CeA reveals six topographically segregated and functionally distinct regions. **A)** Representative CeA coronal sections indicating the areas used for the principal component analysis (PCA). **B-C)** PCA of the average number of c-Fos expressing cells in each sample (average of 3-4 mice per sample) after injections of cocaine, SKF 38393, quinpirole and haloperidol. 2D (**B**) and 3D (**C**) scatter plots show that samples in the same region tend to cluster together. **D)** Schematic of the CeA summarizing the distribution of DAergic fibers and effect of DAergic drugs. CeLf, lateral division of the CeA, frontal part; CeLc, lateral division of the CeA, caudal part; CeCfd, capsular division of the CeA, frontodorsal part; CeCfv, capsular division of the CeA, frontoventral part; CeCc, capsular division of the CeA, caudal part; CeM, medial division of the CeA.

## Discussion

Although DA innervation at the CeA has been largely studied [9,18–20], it remained unclear whether inputs arriving from the vPAG/DR and the VTA/SNc converge into the same CeA regions or rather constitute separate pathways that regulate the CeA with distinct properties. In this work, we demonstrate that the vPAG/DR and the VTA/SNc pathways assemble two different, mostly non-overlapping, circuits (**Fig. 3**). Furthermore, the topographical organization of DA fibers impinging the CeA allowed us to distinguish several regions among the three classical divisions of the CeA, each with specific reactivity to different DA agonists and antagonists (**Fig. 4** and **5**). Together, these results suggest that vPAG/DR and VTA/SNc modulate distinct subsets of topographically segregated CeA neurons which are likely involved in different, although not necessarily unrelated, functions.

Classically, it has been established that DA reuptake by DAT is a fundamental mechanism for terminating DAergic neurotransmission and, accordingly, the presence of DAT has been largely recognized as a marker of DAergic neurons. However, here we have demonstrated that DAergic vPAG/DR neurons have atypically low DAT levels compared to those found in VTA/SNc neurons. Similarly, limited DAT expression has been reported in VTA neurons projecting to the prefrontal cortex, nucleus accumbens medial shell and core and basolateral amygdala [25]. The lower levels of DAT observed in the PAG/DR is likely the consequence of an all or nothing *Dat* expression level in each individual neuron, as we observed in *Dat*^+/I*IRES-Cre*^.Ai14 mice in which only half of TH positive neurons express *Dat*. However, the lower intensity of DAT immunostaining found in the CeL (**Fig. 2A** and **Fig. 3B**) compared with that shown by the reporter *tdTomato* in *Dat*^*+/IRESCre*^.Ai14 mice (**Supp. Fig. 2**) suggests that even neurons with transcriptionally active *Dat* express this gene at much lower levels than canonical DAergic neurons. The functional meaning of low DAT levels in the time-course of DAergic neurotransmission in the CeA should be further investigated in future studies, together with the effects of psychostimulants such as amphetamine and cocaine [26,27].

The comparison of different Cre-driver mouse lines capable of targeting DA neurons is of high interest for a wide community of neuroscientists. In general, DAT-Cre mice have shown to be more selective than TH-Cre mouse lines to target DA neurons in the VTA [28–30] and in the vPAG/DR [31]. However, DAT-Cre lines were reported to have reduced penetrance in the VTA [30] due to reduced *Dat* expression in specific populations of VTA DAergic neurons [25,32]. In agreement with the heterogeneous presence of DAT in VTA neurons, our results also show that DAT is present in ∼50% of DAergic neurons of the vPAG/DR, indicating a limitation of DAT-Cre lines to target DAergic neurons. Of note, given that we used a Cre-reporter mouse line [22] instead of viral vectors-mediated gene delivery, the incomplete penetrance we show in our study cannot be attributed to a partially efficient transduction rate. In addition, we also noticed a population of *Dat* expressing neurons with undetectable TH, which were labeled with retrobeads injected in the CeA (**Fig 2 B-E**). Those DAT+ and TH-neurons were small sized, rounded/oval shaped and located near the aqueduct of Sylvius, as previously described [33]. Of note, neurons with those characteristics have been proposed to be DAergic and to express low *Th* levels, as found in *Th*-GFP and *Pitx3*-GFP mouse lines [34].

The detailed molecular neuroanatomical study performed here sheds light on the intricate organization of the CeA, the complexity of the DAergic innervation that each CeA subdivision receives and their heterogeneous responses to DA. Although the CeA is classically divided in CeC, CeL and CeM, the segregated DAergic innervation and the effect of DA compounds revealed a compartmentalized organization in which three regions in the CeC (frontodorsal, frontoventral and caudal) and two in the CeL (frontal and caudal) are distinguished (**Fig 5D**). This finding is particularly relevant because most functional studies performed to date have focused in caudal regions of the CeA and studied together the CeC and CeL, as if they were a single nucleus. This simplification has limited the study of amygdalar circuits and their implications in emotional behaviors. For example, the role of D2Rs in the CeA seems to be controversial, with reports suggesting both anxiogenic [12,14] as well as anxiolytic [8,16,17] effects. Interestingly, our results show that frontal and caudal CeL display opposite responses to D2R stimulation and blockade (**Fig. 4E, CeL**), suggesting that small variations when placing stereotaxic-guided cannulas into the CeA may lead to different behavioral results, and therefore, seemingly diverging conclusions about the role of D2R in the CeA. Thus, this work highlights the need for more precise approaches to dissect the functional role of regions and particular neuronal populations of the CeA and also calls for a cautious interpretation of the current available data.

## Methods

### Mice husbandry

*Dat*^*+/IRES-Cre*^ [21], Ai14 [22] and *wild-type* mice were maintained in a C57BL/6J background. Mice were housed in ventilated cages under controlled temperature and photoperiod (12 h light/12 h dark cycle, lights on from 7:00 AM to 7:00 PM), with tap water and laboratory chow available *ad libitum*, and separated by sex. All the experiments were performed on mice of both sexes and older than 8 weeks. All procedures followed the Guide for the Care and Use of Laboratory Animals, United States Public Health Services (2011) and were approved by the Institutional Animal Care and Use Committee of INGEBI-CONICET.

### Stereotaxic surgeries

Mice were anesthetized with ketamine (100mg/kg; i.p.) and xylazine hydrochloride (10mg/kg; i.p.). A 10 µl Hamilton syringe connected with a 36-gauge metal needle was used to infuse adeno associated viral vectors or retrobeads using a microsyringe pump at 0.1 µl/min. All stereotaxic coordinates were in relation to the Bregma according to [23]; for the CeA: anterior-posterior, −1.5mm; medial-lateral, +/−3.0mm; dorsal-ventral, −4.9mm; for the VTA/SNc: anterior-posterior, −3.1mm; medial-lateral, +/−0.4mm; dorsal-ventral, −4.5mm; for the PAG/DR: anterior-posterior, −4.1mm; medial-lateral, −1.8mm; dorsal-ventral, −2.1mm, angle: 20° (to avoid the aqueduct). Following infusion, the needle was kept at the injection site for 5 min, and then slowly withdrawn to half way, kept there for 2 more min and then slowly withdrawn outside the brain. Skin was sutured, local anesthesia (lidocaine gel) was applied followed by the analgesic flunixin meglumine (5mg/kg, s.c.). Mice were maintained on a regulated warm pad and monitored until recovery from anesthesia. For retrograde tracing, 0.3 µl of green retrobeads (Lumafluor, Green Retrobeads IX) were injected in a 1:4 dilution of the commercial stock in NaCl 0.9%. For anterograde tracing experiments, pAAV-Ef1a-DIO-hChR2(H134R)-*mCherry*-WPRE-pA, serotype 2 (5.2×10^12^ viral particles/ml, UNC Vector Core) were injected into the VTA/SNc (0.3 µl) or the PAG/DR (1.0 µl) of compound *Dat* ^+/IRES-Cre^ .Ai14 mutant mice.

### Drug administration and c-FOS detection

All drugs were dissolved in NaCl 0.9% to reach a concentration such that the injected i.p. volume was 0.1 ml per 10 g of mouse weight. Experiments evaluating cocaine effects were performed separately from those evaluating quinpirole and haloperidol, and because of that they were not included in the same statistical analysis. Vehicle (NaCl 0.9%), cocaine hydrochloride (20 mg/kg; Sigma), quinpirole (1 mg/kg; Sigma), haloperidol (0.3 mg/kg; Tocris) or SKF 38393 (4 mg/kg; hydrobromide; Tocris) were injected i.p.. Mice were left in their home cage and 90 min later were perfused for tissue fixation and histology.

### Perfusion and histology

Transcardiac perfusion was performed with 5 ml of phosphate buffered saline (PBS, 0.9% NaCl, 2.7 mM KCl, 10 mM K2HPO4, 2 mM KH2PO4, pH 7.5) followed by 50 ml of paraformaldehyde 4% in PBS and brains were removed and post-fixed in the same solution at 4°C for 12-16 h. Brains were sectioned at 40 µm on a vibratome (Leica) and used immediately or stored at −20°C in a solution containing 30% (v/v) ethylene glycol, 30% (v/v) glycerol and PBS, until they were processed for immunofluorescence. Immunolabeling was performed as follows: free-floating sections were rinsed three times for 10 min in PBS and then incubated for 16 h at 4°C in primary antibody solution with normal goat serum 2% (w/v), Triton X-100 0.3%, in PBS. The following primary antibodies were used: rabbit anti-TH (1:2,000; Millipore, AB5935), chicken anti-TH (1:1,000; Abcam, AB76442), rabbit anti-C-FOS (1:500, Santa Cruz Biotechnology, SC-52), rabbit anti-dsRed (1:500; Clontech, 632496), rat anti-DAT (1:500, Millipore, MAB369). After incubation with a primary antibody, sections were rinsed twice for 20 min in PBS and then incubated for 2 h at room temperature with goat or donkey Alexa-488 or Alexa 555 coupled secondary antibody 1:1,000 in Triton X-100 0.3% in PBS. Finally, sections were rinsed twice for 20 min in PBS and mounted with Vectashield (Vector Labs) for confocal microscopy or glycerol 50% (v/v) in PBS for fluorescence and bright field microscopy.

### Microscopy and images analysis

Confocal images for co-expression analysis (**Fig. 2**) were obtained using a Leica Confocal TCS-SPE microscope. Co-expression was manually quantified using the tool “Cell Counter”. Images for c-FOS quantification by anatomical region, innervation analysis, DAT and TH intensities measurements and anterograde tracing were obtained by fluorescence microscopy. Images were analyzed with the Fiji platform [43] of the ImageJ software [44]. Coronal sections from AP: −0.94 to AP: −1.06 mm for frontal regions of the CeA and sections from AP: −1.22 to AP: −1.58 mm for caudal regions of the CeA were analyzed.

Analysis of TH and DAT intensities and DAT/TH ratios **(Fig. 1, Fig. 3B, Suppl. Table 1**) were obtained as follows: regions from CeA, dorsal striatum and medial amygdaloid nucleus posterodorsal (MePD) were delimitated and average intensities for TH and DAT channels were calculated using the tool “Measure”. MePD was found to have considerable low signal for TH and DAT and was considered as background signal. For each section, MePD signal (background) was subtracted from values of other areas and then values from each CeA region was relativized to dorsal striatum values. Data of the same region from different slices were averaged to obtain intensity values shown in **Fig. 1** and **Suppl. Table 1**. For each slice, normalized DAT values were divided by normalized TH values, and DAT/TH values of the same region from different sections were averaged to obtain DAT/TH ratios showed in **Suppl. Table 1**.

For **Fig. 4B,D**, cell number was semi-automatically quantified with the tool “Analyze particles” after manually delimitating the region of interest. For statistics, number of cells in each nucleus was obtained adding the values of the corresponding nucleus for each of all sections, and the number of sections added was later used as offset in the Statistical analysis. For graphs, the mean of cells between sections was calculated for each nucleus.

For **Fig. 4E** and **Fig. 5**, images were automatically processed using Python with the libraries Numpy and SciPy. Between three to four coronal sections of each mouse, corresponding to AP 0.94, 1.06, 1.34 and 1.58 mm from Bregma, were analyzed (**Fig. 5A**). For each coronal section of every mouse, images were read as Numpy and transformed to binary such that only vales above the 95.5 percentile were 1 and any other value was 0. Two squared samples where delimited in each region, and for each sample particles above the threshold where identified and labeled and particles of size above 30 pixels where quantified. Importantly, the location of the samples at each coronal section were the same for every mouse, which allowed to calculate the average number of cFos positive nuclei across mice for each drug and sample to perform the principal component analysis with the pharmacological profile of each sample (**Fig. 5B-C**). For **Fig. 5D**, the average between samples of each region was first calculated for each drug and mouse to avoid, and then the average and 95% confidence interval across mice for each drug was plotted.

### Statistical analysis

All data represent the mean ± 95%CI and were graphed using Python (libraries matplotlib and seaborn). Statistical analysis were performed in Python (packages numpy, pingouin, statsmodels and sklearn) for **Figs. 1, 2 and 5** or R Studio (libraries glmmTMB and lsmeans) for **Fig. 4**. The statistical analysis are indicated in the Figure legends. Discrete data of **Fig. 4B and D** were analyzed with generalized linear mixed model (GLMM) with Poisson error structure (link:loggit) or Negative-binomial error structure when data had overdispersion. Assumptions for the GLMM were evaluated assessing the absence of patterns in Pearson’s residuals graph and calculating the dispersion parameter to assess subdispersion or overdispersion. The principal component (**Fig. 5B-C**) analysis was performed using the average number of cFos positive nuclei of each sample between mice for each drug (columns) and sample (rows). Principal components across drugs were calculated and used to plot scatter plots of each sample.

## Code and accessibility

All data presented in this paper and codes for plotting and statistical analysis can be accessed at https://github.com/casey-e/Casey-et-al-2022-b.

## Conflict of Interest

The authors declare no competing financial interests.

## Acknowledgments

This work was supported by grants from the Agencia Nacional de Promoción Científica y Tecnológica, Argentina (M.R.) and a doctoral fellowship from Consejo Nacional de Investigaciones Científicas y Técnicas (CONICET), Argentina (E.C.).

## Notes

https://github.com/casey-e/Casey-et-al-2022-b

### Competing Interest Statement

The authors have declared no competing interest.

