## Supplementary Figures and Tables for "Dopaminergic innervation at the central nucleus of the amygdala reveals distinct topographically and functionally segregated regions"

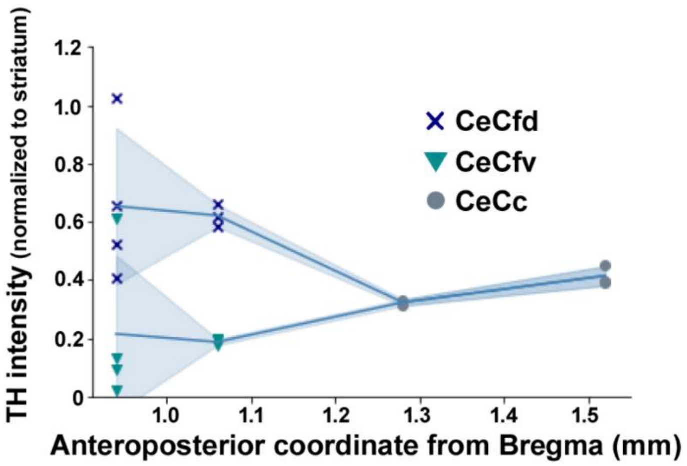


**Suppl. Fig. 1. TH gradients in the CeC.** Quantification of TH intensity in the CeC, across the fronto-caudal axis, differentiating among fronto-dorsal, fronto-ventral and caudal regions. Values are normalized to intensity in the dorsal striatum of the same coronal section.

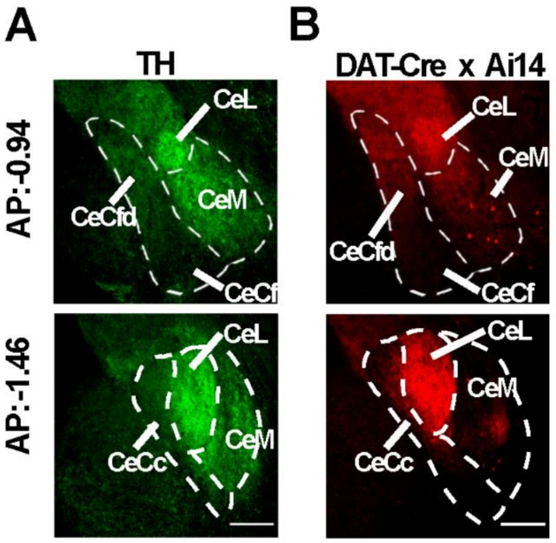


**Suppl. Fig. 2. Identical distribution of TH immunostained fibers and genetically marked DAT expressing fibers.** Representative histology of the CeA of a *wild-type* mouse immunostained against TH (**A**) and the CeA of a *Dat*$^{+/IRES\text{-}Cre}$.Ai14 mouse (**B**). Upper row: frontal part of the CeA, from AP:-0.82 to AP:-1.06 mm (the example corresponds to AP:-0.94 mm); bottom row: caudal part of the CeA, from AP:-1.22 to AP:-1.70 (the example corresponds to AP:-1.46 mm).

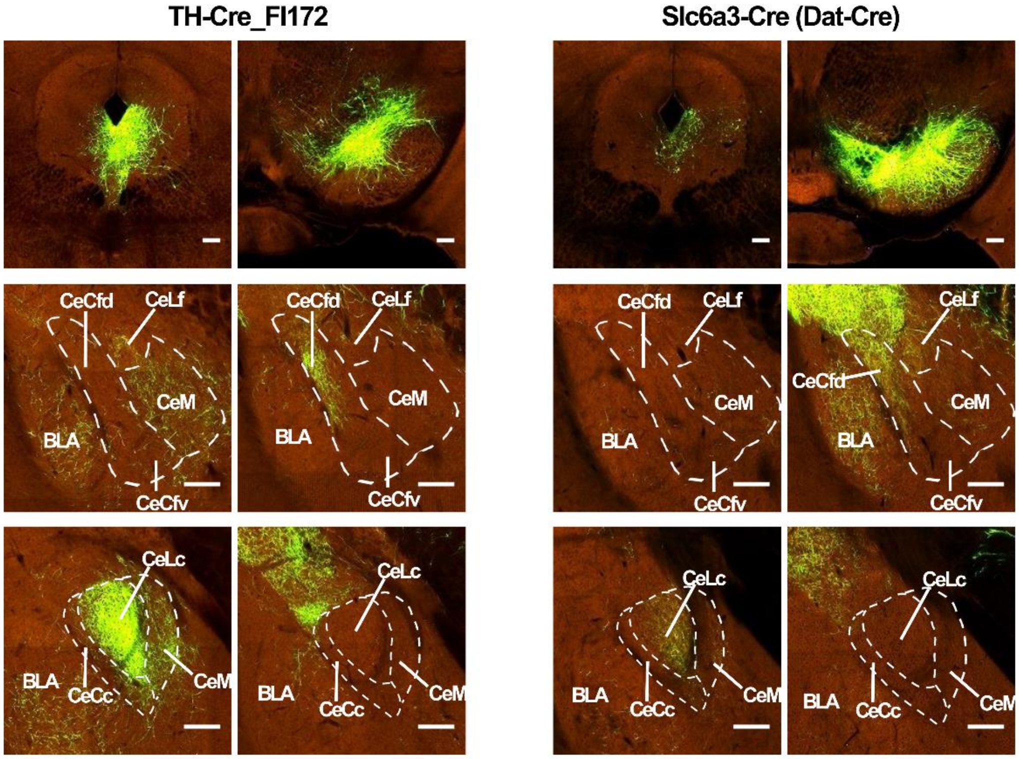


**Suppl. Fig. 3. PAG/DR DAergic neurons project to CeL and CeM, and VTA/SNc DAergic neurons project to frontodorsal CeC.** Data from mouse connectivity of Allen Brain Atlas (http://www.brain-map.org/) corresponding to TH-Cre (TH-Cre_FI172) and *Dat*-Cre (Slc6a3-Cre) mice injected with a Cre-dependent anterograde marker in the vPAG/DR or the VTA/SNc. Top row, injection sites; middle row, frontal CeA (AP:-1.06 mm); low row, caudal CeA (AP: -1.46 mm). TH-Cre and *Dat*-Cre mice show identical qualitative innervation patterns with higher intensities in the TH-Cre line. BLA, basolateral amygdala; CeLf, lateral division of the CeA, frontal part; CeLc, lateral division of the CeA, caudal part; CeCfd, capsular division of the CeA, frontodorsal part; CeCfv, capsular division of the CeA, frontoventral part; CeCc, capsular division of the CeA, caudal part; CeM, medial division of the CeA.

**Suppl. Table 1.**

| CeA division | Normalized TH intensity | Normalized DAT intensity | DAT/TH |
|---|---|---|---|
| **CeCfd** | 0,6 (0,61-0,58) | 0,32 (0,35-0,29) | 0.54 (0.58-0.5) |
| **CeCfv** | 0.17 (0.17-0.17) | 0.02 (0.02-0.02) | 0.11 (0.08-0.15) |
| **CeCc** | 0.34 (0.36-0.33) | 0.05 (0.06-0.05) | 0.15 (0.16-0.14) |
| **CeLf** | 1,59 (1,52-1,65) | 0,52 (0,56-0,48) | 0.32 (0.36-0.29) |
| **CeLc** | 1.14 (1.21-1-07) | 0.16 (0.18-0.14) | 0.14 (0.16-0.13) |
| **CeM** | 0.75 (0.84-0.66) | 0.09 (0.11-0.08) | 0.12 (0.13-0.12) |

Intensities of fluorescence of TH and DAT immunolabeling in each CeA division were normalized to intensity of fluorescence of the dorsal striatum in the same histological section and the ratio between DAT and TH normalized intensities were calculated. CeLf, lateral division of the CeA, frontal part; CeLc, lateral division of the CeA, caudal part; CeCfd, capsular division of the CeA, frontodorsal part; CeCfv, capsular division of the CeA, frontoventral part; CeCc, capsular division of the CeA, caudal part; CeM, medial division of the CeA. Mean ± individual data is shown (n=2 mice).

**Suppl. Table 2.**

| Principal component | Explained variance | Cocaine contribution | SKF 38393 contribution | Quinpirole contribution | Haloperidol contribution |
|---|---|---|---|---|---|
| **PC1** | 0.51 | -0.16 | -0.12 | -0.16 | 0.96 |
| **PC2** | 0.25 | 0.34 | -0.00 | 0.90 | 0.21 |
| **PC3** | 0.14 | 0.18 | 0.96 | -0.10 | 0.13 |

Explained variance of each component and contributions of the original variables.
